# Drivers of antibiotic resistance in two monsoon-impacted Indian urban rivers receiving untreated wastewater

**DOI:** 10.1101/2024.10.31.621109

**Authors:** Kenyum Bagra, Harshita Singh, Uli Klümper, Gargi Singh

**Author notes:** Corresponding authors; Dr. Gargi Singh, Department of Civil Engineering, Indian Institute of Technology Roorkee, Uttarakhand, India, Dr. Uli Klümper, Institute of Hydrobiology, TU Dresden, Dresden, Germany.

## Abstract

Rivers in India receive 78.7% of untreated sewage from cities and towns making it a potential global “hotspot” for AMR. During monsoon, flooding is common in most Indian cities bringing in fecal contamination in storm runoff. Identifying the major driving factors contributing to the elevated levels of AMR in these rivers is essential to combat AMR proliferation. Determining whether the introduction of ARBs and ARGs through fecal pollution or subsequent (co-) selection for ARGs through chemical pollutants is responsible for the rise and spread of AMR is important in mitigation efforts. To achieve this, we targeted two rivers in northern India which received untreated wastewater and assessed the level of two fecal indicators (*E. coli* and *int*I1) and wastewater-associated ARGs (*erm*F, *sul*1, *sul*2, and *tet*W) along with heavy metals (Ag, Cd, Co, Cr, Cu, Fe, Mn, Ni, Pb, and Zn). Water and sediment samples were collected in triplicates at 5-6 locations for each river. Consequently, to evaluate how seasonality affects the ARG levels as well as their drivers in these heavily polluted rivers, sampling campaigns were carried out during three seasons: in summer (pre-monsoon), during monsoon, and post-monsoon in winter. We here demonstrate that the main source for the river resistome in the Bindal and Rispana is the constant inflow of untreated wastewater throughout the year and not the co-selection of ARGs due to the presence of heavy metals. This may be true for many Indian rivers that receive untreated or partially treated wastewater. The rainfall adds to the ARG abundance of the river instead of diluting it. MGE *int*I1 is more reliable than *E. coli* as an indicator for fecal pollution and horizontal gene transfer. Heavy metals, despite being present in the river in higher concentrations than the MCSC, no evidence for co-selection was observed.

## Introduction

The rise of antimicrobial resistance (AMR) has been considered a silent pandemic on the global scale [1]. Based on global clinical reports, 4.95 million people succumbed to bacterial infections with resistant pathogens in 2019 [2], with numbers expected to rise up to 10 million deaths per year by 2050 [3]. As the threat of AMR is growing, there is a need to stop the widespread proliferation of AMR not only in clinical settings but also due to environmental contamination with resistant bacteria and selective/co-selective agents [4], [5].

The aquatic environments are not only predominant reservoirs but also are the predominant conduits of circulation of antibiotic-resistant bacteria and antibiotic-resistance genes (ARGs) in the environment and drivers of subsequent exposure to communities. The discharge of wastewater from municipal, agricultural, and industrial sectors into rivers is a well-established pathway for AMR and selectors/co-selectors to enter the environment [6]. It also makes rivers a suitable breeding ground for AMR and a source of AMR contamination and human and animal exposure [7]. Raw sewage contains pollutants like heavy metals, disinfectants, biocides, pharmaceuticals, and antibiotics, along with the gut microflora of humans and animals (including pathogens) enriched in carried antibiotic-resistant genes (ARGs) [8], [9]. Even with proper wastewater treatment, these emerging contaminants are not completely removed, resulting in many studies documenting elevated levels in rivers after WWTP discharge points [10], [11]. The simultaneously released (co-) selecting chemicals can further induce selective pressure on the river microbial community, resulting in increased acquisition and selection rates of ARGs [12], [13]. Heavy metals are among the most relevant environmental contaminants with co-selection potential for AMR in diverse environments, including rivers [13], [14].

The easy access to antibiotics without prescription has resulted in India becoming the highest global consumer of antibiotics [15]. Thus, the prevalence of AMR at the community level in India is very high [15]. A study had reported that 12 out of 18 Swedish exchange students visiting various Indian cities upon return acquired ESBL-producing *E. coli* [16]. Moreover, the student’s gut microbiomes showed increased resistance to sulphonamide, trimethoprim, beta-lactams tetracycline and increased integrases and insertion sequence common regions elements (IS*CRs*) responsible for HGT. As guts are heavily colonized among Indian citizens, high loads of AMR bacteria and their ARGs are released into wastewater [17]. India, however, has a limited WWTP infrastructure, resulting in almost 78.7% of its sewage from cities and towns entering rivers untreated [18]. Indian rivers are, hence, a potential global “hotspot” for AMR. Six years after the first documented presence of *bla*_NDM-1_ in 2008, a study reported *bla*_NDM-1_ present during a massive influx of seasonal pilgrims visiting the upstream of the holy Ganga River [19]. A metagenomic study of sediments from the Mathua River in Pune found that the carbapenemase-resistant gene (*bla*_OXA-58_) positively correlated with the pathogen *Acinetobacter* present [20]. The same study also detected *mcr*-1, a gene conferring resistance to macrolides, e.g., Colistin, a last resort antibiotic, in the river sediment and upstream river, despite it being recognized as a relatively pristine area. In addition to contamination with fecal bacteria and their ARGs, Indian rivers also receive significant amounts of other chemical pollutants, including diverse heavy metals, which are commonly known as potent co-selective agents for ARGs [21], [22], [23], [24]. Elevated levels of ARGs and these co-selectors have been simultaneously detected in a vast number of Indian rivers [20], [24], [25], with Reddy et al. reporting a strong correlation between metal resistance and ARGs in the river Ganges [24].

In this context, heavily polluted rivers used as conduits for raw urban sewage and other urban waste present a unique yet under-investigated environment for AMR proliferation. One such case is the sub-tributaries of Ganga in Dehradun: the Bindal and Rispana receive not only ∼30% untreated wastewater [26] but solid waste, leading to elevated levels of microplastics and heavy metals in the rivers [27]. Identifying the major driving factors contributing to the elevated levels of AMR in these rivers is essential to combat AMR proliferation. Here, especially the question if introduction through fecal pollution or subsequent (co-)selection for ARGs through chemical pollutants are responsible and should mainly be targeted in mitigation efforts remains to be answered. To achieve this, we targeted two rivers in northern India which received untreated wastewater and assessed the level of two fecal indicators (*E. coli* and *int*I1) and wastewater-associated ARGs (*erm*F, *sul*1, *sul*2 and *tet*W) along with heavy metals (Ag, Cd, Co, Cr, Cu, Fe, Mn, Ni, Pb, and Zn). Water and sediment samples were collected in triplicates at 5-6 different locations for each river. Moreover, both river tributaries are heavily influenced by seasonality. Dehradun, similar to other regions in India, undergoes a monsoon season characterized by heavy rainfall from June to September. During the peak monsoon period in August, the rainfall intensifies, often leading to flooding, particularly in low-lying areas at the peak of the monsoon. Studies reported high levels of fecal contamination in storm runoff correlating increased ARG levels [8], [28], [29]. Such heavy rainfall events have been shown to come with contradictory effects on ARG levels. While rises in highly urbanized areas were detected, the incoming water can also dilute ARG abundances in more pristine areas [30]. Consequently, to evaluate how seasonality affects the ARG levels as well as their drivers in these heavily polluted rivers, sampling campaigns were carried out during three seasons: in summer (pre-monsoon), during monsoon, and post-monsoon in winter.

## Material & Methodology

### Study location and sample collection

Water and sediment samples were collected from multiple locations (SI 1) on the banks of two sub-tributaries of the river Ganga – the Bindal and Rispana, flowing through Dehradun, the capital city of Uttarakhand, India. Dehradun is highly urbanized, with many slum areas packed along the stretch of the river and stretches to an area of 300 km^2^ (29°58′ to 31°25′ N, 77°34′ to 78°18′ E). The monsoon period in Dehradun occurs between June and September, with the peak of the monsoon precipitation level reaching up to an average of 715 mm in the month of August [31]. The Bindal and Rispana flow parallelly through Dehradun and merge to form the Suswa River. Suswa then merges with the river Song and meets Ganga further downstream near Rajaji National Park. The Bindal is a seasonal river that primarily acts as an urban drain for the city for most of the year, as it receives roughly 18.14 MLD municipal treated wastewater from an WWTP [26] and from urban stormwater drains that frequently carry both stormwater and greywater along with the solid waste. Rispana is the city’s primary water source for irrigation and groundwater recharge. It receives around 9.386 MLD of municipal wastewater [26] and also greywater from the urban drains.

The sampling locations B1 and R1 are in a relatively less densely populated area upstream of Dehradun city on the Bindal and Rispana, respectively. Before and after the monsoon, the Bindal was dry upstream to location B1. Some of the sampling sites were situated near slum areas, such as B2 and R1. Samples were also collected upstream and downstream of the WWTPs discharging into the Bindal and Rispana, with sampling points designated as B3 (upstream) - B3D (downstream) and R4 (upstream) - R4D (downstream), respectively. The last sampling points, B4 and R5, are situated just before the rivers’ confluence into Suswa river. The two rivers join downstream into another river named Song, which then meets Ganga further downstream. Samples were collected in July, August, and November of 2019 in triplicates for biological analysis in a sterile 50 mL centrifuge tube. Water samples were collected in 50 mL amber bottles for metal analysis, and the sediment samples were collected in 50 mL sterile centrifuge tubes.

### Molecular biological analyses

#### DNA extraction and qPCR

The water samples were aseptically filtered on 0.22 µm mixed cellulose ester filter papers (Axiva Achichem Biotech, Delhi, India), which were then used to extract DNA. The Qiagen DNeasy® Soil extraction kit (Qiagen®, Hilden, Germany) was used to extract DNA from the residue on the filter paper and sediments following the manufacturer’s protocol. The DNA was eluted in 100 μL of molecular biology-grade water and stored at -20 °C until further analysis. The samples were screened for the presence/absence of specific ARGs, MRGs, fecal indicators, and pathogens using quantitative polymerase chain reaction (qPCR) on Prima-96™ Thermal Cycler (HiMedia Laboratories Pvt Limited, Mumbai, India). ARGs-*ermF*, *sul*1*, sul*2*, tet*W and class 1 integron-integrase gene cassette, *intI*1, and fecal indicator *E. coli* were detected by gel electrophoresis using Hi-Gel run1014 (HiMedia Laboratories Pvt Limited, Mumbai, India) and were quantified using qPCR (Primer list, Table AI2, Annexure-I). Each qPCR reaction consisted of 5 μL of SYBR Green, 2 μL of forward and reverse primers and 1 μL of DNA template. Each qPCR run included a standard curve covering eight orders of magnitude of standard and three negative controls that used molecular biology-grade water as a DNA control. The quality and concentration of the extracted DNA were determined by a Nanodrop OneC spectrophotometer (Thermo Scientific™, Massachusetts, USA). The qPCR standards were prepared by cloning the target amplicons on TOP10 competent cells using the TOPO TA Cloning kit (Invitrogen, CA, USA). Each qPCR run included a standard curve covering eight orders of magnitude and a melt curve analysis with a temperature gradient from 50 °C to 95 °C at the end to confirm the amplified products. The cycling conditions consisted of activation at 50 °C for 2 min, initial denaturation at 95 °C for 2 min, and 40 cycles of denaturation (95 °C for 15 s) followed by combined annealing and extension (60 °C for 1 min). When the Ct value fell below the limit of quantification (LOQ), the data was considered censored [32]. To minimize potential biases, the Ct values below LOQ were imputed with the value of LOQ/2 for the corresponding qPCR assay if the censored data was less than 40% [33], [34].

### Metal analysis

Heavy metals Ag, Cd, Co, Cr, Cu, Fe, Mn, Ni, Pb, and Zn were analysed for water and sediment samples using inductively coupled plasma mass spectrometry (ICP-MS) (Agilent 7900, Agilent Technologies, Inc, USA). For sample preparation, 2 mL HNO_3_ and 2 mL H_2_O_2_ were added to a 10 mL water sample and 1 g sediment. The resulting mixture was diluted to 100 mL and digested using a hot plate until the volume was reduced to 20 mL. Sediment samples were dried at 105 ℃ for 24 hours in a hot oven (AI-7981, i-therm, Maharashtra, India) and homogenized before digestion. After digestion, the samples were filtered and diluted at 1:10 ratio for water samples and 1:100 for sediment samples before analysis.

### Statistical analysis

The statistical analysis was performed in the R version using RStudio v4.1. as a GUI [35]. The Kruskal-Walli’s test was employed to evaluate differences in fecal indicators and ARGs among different locations due to the non-parametric nature of the data. This test was followed by Dunn’s post-hoc test to identify specific location-wise differences. Spearman’s rank correlation coefficient was used to assess the relationship between fecal indicators and ARGs, as well as between 16S rRNA gene copies and heavy metal concentrations. The Wilcoxon signed-rank test was applied to analyse seasonal variations in fecal indicators and ARGs at each location. To analyse seasonal variation in fecal indicators and ARGs at each location, the Wilcoxon signed-rank test was applied. This test is suitable for comparing paired samples and detecting differences between seasonal time points [36].

We used a linear mixed-effects model to assess the effects of season (categorical variable) and sample location based on river flow direction (continuous variable) on microbial loads. The model included season as a fixed effect and flow direction as a continuous predictor. The model also accounted for repeated measurements at different locations by including location as a random effect. The analysis was conducted separately for each dataset (e.g., Bindal Water, Bindal Sediment, Rispana Water, Rispana Sediment). Model parameters were estimated using Restricted Maximum Likelihood (REML), and interaction terms were included to explore how the effect of season varied along the river flow direction. The linear mixed-effects model used in the analysis can be described by the following formula:

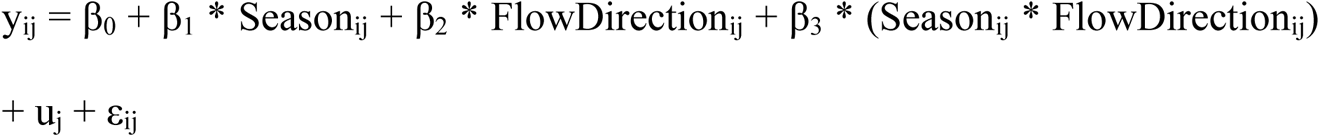

Where y_ij_ is the target abundance for the i-th observation at the j-th location, β_0_ is the intercept (the baseline microbial load), β_1_, β_2_, and β_3_ are the fixed-effect coefficients for season, flow direction, and their interaction, u_j_ is the random effect for location (to account for variability between locations) and ε_ij_ is the residual error term for the i-th observation at the j-th location.3.

## Results

### Bacterial loads in the Bindal and Rispana rivers

First, to assess the total bacterial load dynamics in the two Indian urban rivers - Bindal and Rispana - the 16S rRNA gene was quantified. The total bacterial count of the river water samples based on the 16S rRNA gene varied between seasons at the different locations on the Bindal river: summer: 6.56 ± 0.28 to 7.43 ± 0.43 log_10_ copies mL^-1^; monsoon: 5.58 ± 0.16 to 7.01 ± 0.45 and winter: 8.1 ± 0.04 to 8.53 ± 0.04) and Rispana river (summer: 5.37 ± 0.13 to 6.82 ± 0.15 log_10_ copies mL^-1^; monsoon: 5.2 ± 0.02 to 7.92 ± 0.14; winter: 6.6 ± 0.07 to 8.44 ± 0.12; Fig. 1). A linear mixed-effects model (fit with REML) was used to analyze the effect of season (fixed effect) and sample locations based on river flow direction (continuous variable) on total bacterial loads in water samples from each river. The model revealed a significant increase in total bacterial loads during the Winter compared to the monsoon (Bindal: p = 0.024, ηp² = 0.52; Rispana: p < 0.001, ηp² = 0.89) and compared to summer (Bindal: p = 0.024; Rispana: p < 0.001). However, no significant difference was found between summer and monsoon (Bindal: p = 0.512; Rispana: p = 0.512). Moreover, there was no significant effect of sample locations based on flow direction or the interaction between season and location. Most interestingly, no effect of wastewater influx on the bacterial load was detected when comparing the locations immediately upstream and downstream of the wastewater treatment plant in each river: For the Bindal, no significant difference in bacterial load of locations B3 and B3D was apparent for any of the seasons (all p > 0.05, Kruskal-Wallis with Dunn’s test), while for the Rispana similarly no effect was detected when comparing locations R4 and R4D (all p > 0.05; Fig. 1). However, when analysing the sediment samples based on the linear mixed-effects model no significant differences in total bacterial loads between seasons were observed (Bindal: p > 0.05, ηp² = 0.39; Rispana: p > 0.05, ηp² = 0.34). Again, flow direction and the interaction between season and flow direction were not significant predictors in either dataset (all p > 0.05). Despite the missing seasonality in the sediment compared to the water datasets, the bacterial load of the water samples and sediment samples across locations and seasons were highly correlated for both the Bindal (p = 5.07 x 10^-5^, R = 0.57; Spearman Rank Correlation) and the Rispana (p = 1.64 x 10^-5^, R = 0.55). The total bacterial count of the river water samples based on the 16S rRNA gene was varied between seasons at the different locations on the Bindal river: summer: 7.69 ± 0.14 to 8.99 ± 0.14 log_10_ copies mL^-1^; monsoon: 8.02 ± 0.3 to 8.53 ± 0.18 and winter: 8.83 ± 0.23 to 9.52 ± 0.24) and Rispana river (summer: 7.47 ± 0.17 to 9.03 ± 0.22 log_10_ copies mL^-1^; monsoon: 7.21 ± 0.6 to 8.71 ± 0.07; winter: 7.74 ± 0.88 to 9.14 ± 0.1; Fig. 1).

**Fig. 1.**
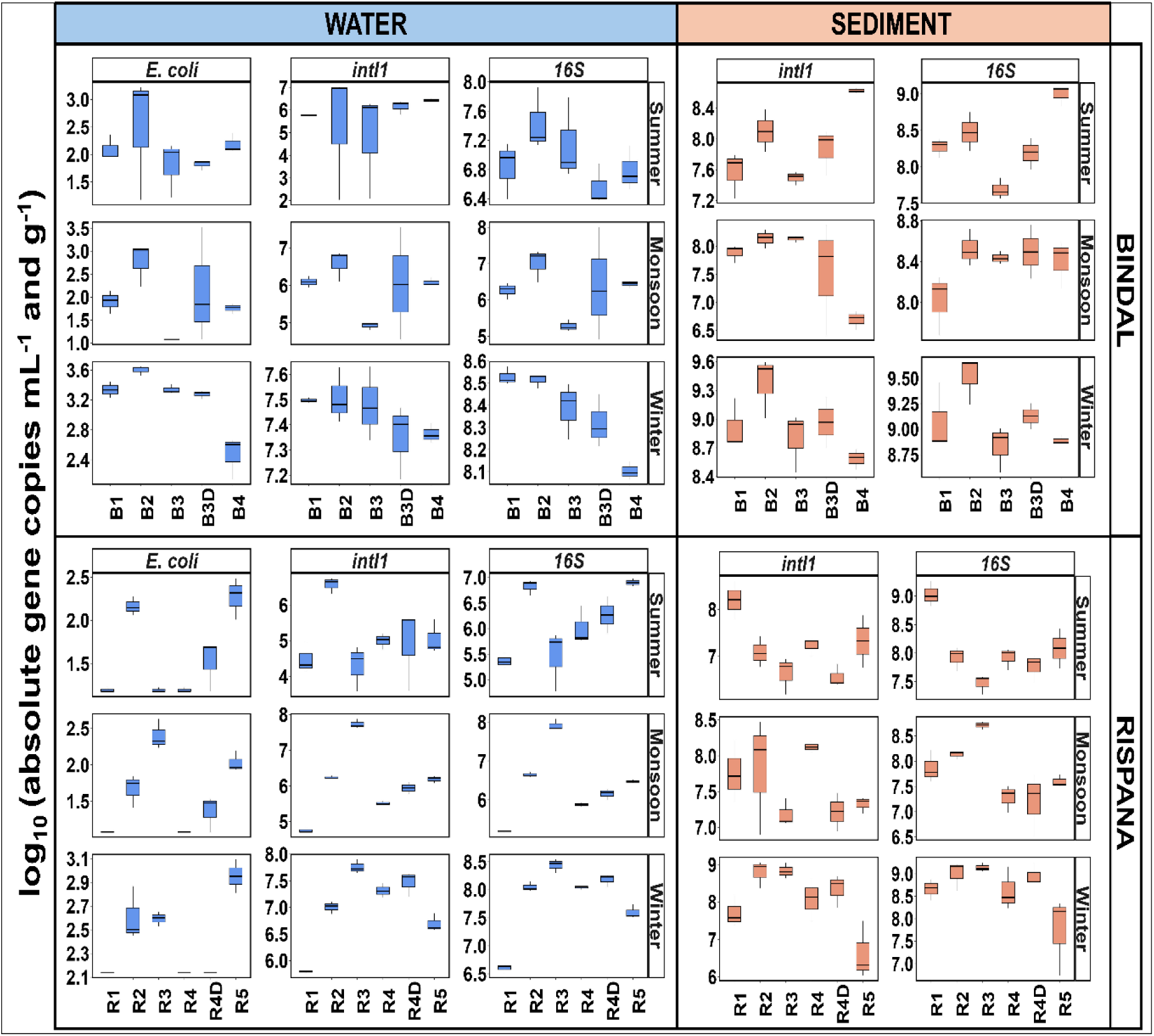
Fecal indicators (E. coli (yccT) and intI1) and total bacteria count (16S rRNA) in water and sediment samples for Bindal and Rispana across the length of the rivers during different seasons (summer, monsoon, and winter). Absolute gene abundance of fecal indicators with total bacteria in water and sediment samples for Bindal and Rispana across the length of the rivers during different seasons.

### Fecal pollution in the Bindal and Rispana rivers

As levels of AMR in rivers are regularly connected to fecal pollution through via wastewater inflow or agricultural field runoff, we analysed the fecal pollution in both rivers at different sampling locations during the three seasons via abundance of two established indicators, *E. coli* (based on the *ycc*T gene) and the class 1 integron integrase gene cassette, *int*I1. The absolute abundance of *E. coli* in the river water samples at each timepoint was highly variable between the different locations of Bindal (summer: 1.81 ± 0.51 to 2.5 ± 1.14 log_10_ copies mL^-1^; monsoon: 1.08 ± 0.51 to 2.77 ± 0.47; winter: 2.47 ± 0.29 to 3.6 ± 0.06) and Rispana river (summer: 1.19 ± 0.03 to 2.27 ± 0.24 log_10_ copies mL^-1^; monsoon: 1.08 ± 0 to 2.4 ± 0.21; winter: 2.14 ± 0 to 2.61 ± 0.23; Fig. 1). While *E. coli* was detectable in all water and sediment samples, it was consistently below the limit of quantification. Similarly, the absolute abundance of *int*I1 in water samples was highly variable across sampling locations in both, Bindal (summer: 4.82 ± 2.37 to 6.43 ± 0.08 log_10_ copies mL^-1^; monsoon: 4.92 ± 0.1 to 6.59 ± 0.42; winter: 7.35 ± 0.15 to 7.51 ± 0.11) and Rispana (summer: 4.29 ± 0.64 to 6.57 ± 0.23, monsoon: 4.75 ± 0.07 to 7.73 ± 0.13, winter: 5.8 ± 0.02 to 7.76 ± 0.13; Fig. 1). Unlike *E. coli*, *int*I1 was detectable in sediment samples for all locations and the abundance was again highly variable in Bindal (summer: 7.5 ± 0.09 to 8.63 ± 0.03 log_10_ copies g^-1^, monsoon: 6.69 ± 0.16 to 8.14 ± 0.17, and winter: 8.59 ± 0.11 to 9.38 ± 0.32) and Rispana (summer: 6.56 ± 0.24 to 8.2 ± 0.4 log_10_ copies g^-1^, monsoon: 7.19 ± 0.19 to 8.12 ± 0.07, and winter: 6.62 ± 0.78 to 8.79 ± 0.36; Fig. 1).

While the absolute abundances of *E. coli* and *int*I1 were highly variable across locations, both fecal indicators were highly correlated with one another (Bindal: R = 0.93, p <0.00001; Rispana: R = 0.74, p = 1.52 x 10^-01^). Similarly, both were strongly positively correlated with 16S rRNA gene abundance (Fig. 1): For Bindal water, 16S rRNA gene copies positively correlated with absolute *E. coli* (R = 0.95, p < 0.0001, Spearman rank correlation) and absolute *int*I1 abundance (R = 0.89, p < 0.0001). Similarly, for Rispana water, the 16S rRNA gene copies positively correlated with *E. coli* (R = 0.87, p < 0.0001) and *int*I1 abundance (R = 0.88, p < 0.001). 16S rRNA gene copies in sediment samples also positively correlated with *int*I1 in Bindal (R = 0.82, p = 3.2 x 10^-12^) and Rispana (R = 0.56, p = 9.588 x 10^-06^). When consequently evaluating the relative abundance of either fecal indicator, no significant changes along the flow of Bindal and Rispana were observed for each season based on the linear mixed-effects model (all p > 0.05; Fig. 2). Surprisingly, in either river, again no effect of wastewater influent on fecal indicator abundance was observed when comparing the up - and downstream the WWTP locations for either of the seasons (all p > 0.05, Kruskal-Wallis with Dunn’s post-hoc test). Hence, in each season, only the total bacterial load changed across sampling locations, while the relative proportion of fecal indicators in the entire community remained relatively stable (Fig. 2). However, between the different seasons, clear differences in the relative fecal indicator load were observed: When analysing the transition from summer to monsoon season, a clear and consistent increase in *int*I1 relative abundance during monsoon across sampling locations was observed in the river samples (p = 0.0185, Wilcoxon signed-rank test; Fig. 3). Similar increases in the relative abundance of *E. coli* by up to 1 order of magnitude higher in monsoon compared to summer were observed, but this trend was not consistent enough across samples for statistical significance (p = 0.7; Fig. 3). After the rainfall event, the relative abundance of fecal indicators dropped significantly in winter for most river locations (*E. coli*: p = 0.00097; *int*I1: p = 0.00097; Fig. 3). However, these trends were not mirrored in the sediment samples, with no significant difference between summer and monsoon (p = 0.0185) or monsoon and winter (p = 0.63; Fig. 3).

**Fig. 2.**
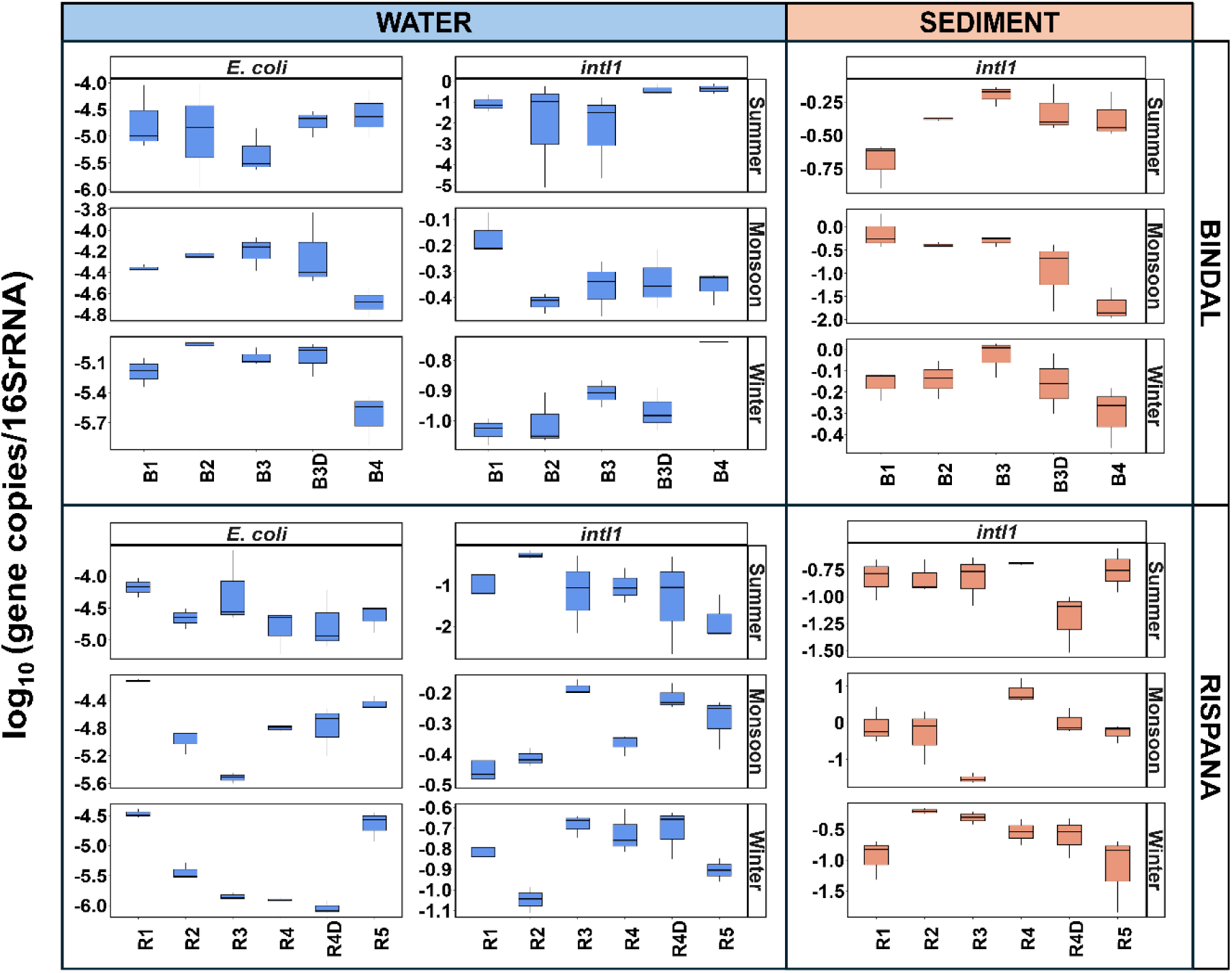
Fecal indicators (E. coli (yccT) and intI1) and total bacteria count (16S rRNA) in water and sediment samples for Bindal and Rispana across the length of the rivers during different seasons (summer, monsoon, and winter). Relative gene abundance of fecal indicators in water and sediment samples for Bindal and Rispana across the length of the rivers during different seasons. The box represents the median with the lower and upper quartile range. Datapoints outside the box are considered outliers.

**Fig. 3.**
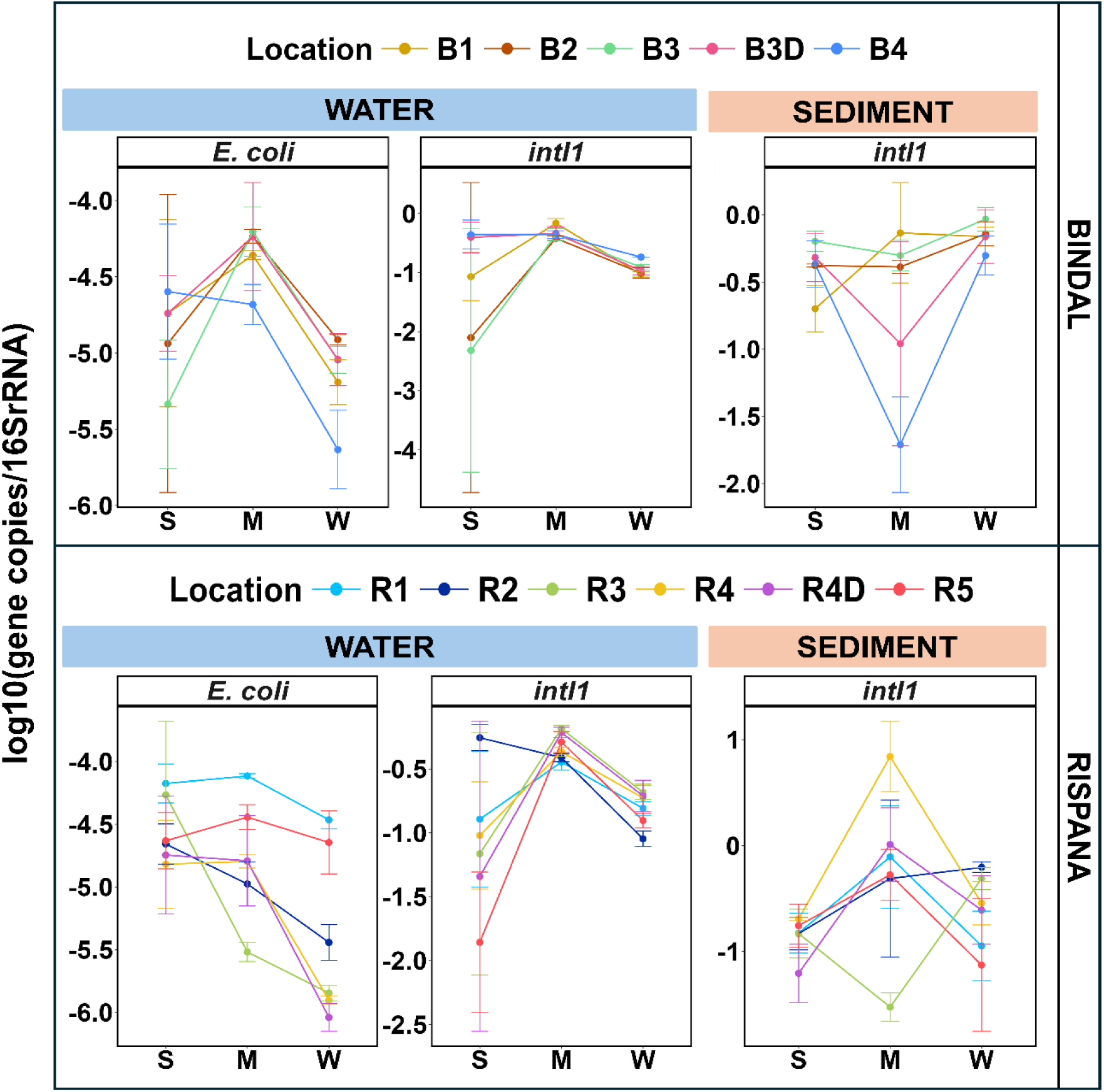
Seasonal variation of the relative abundance of fecal indicators: *E. coli* (yccT) and *int*I1 (gene copy/16S rRNA per mL^-1^) in water and sediment samples for each location in Bindal and Rispana, where B3D and R4D are downstream the WWTP discharge point. The point indicates the mean of the relative gene abundance in the site with standard deviation (n =3), indicating the change with the connecting line across summer, monsoon, and winter for all the samples. *E. coli* (yccT) in sediment samples were below the limit of quantification (LOQ).

In summary, the high absolute presence of *E. coli* and *int*I1 in both rivers indicates strong fecal pollution throughout. However, differences in absolute abundance in certain sites can be explained solely by changes in total bacterial load. No spatial differences between sampling locations in the relative abundance of fecal indicators were observed for either river. Still, significant temporal increases during the monsoon season indicate that rainfall affects the level of fecal pollution in the rivers.

### Heavy metal pollution in Bindal and Rispana rivers

Since in addition to fecal pollution the presence of co-selective agents such as heavy metals that are highly prevalent in urban Indian rivers plays a major role in the spread of AMR, we analysed the heavy metal levels (Ag, Cd, Co, Cr, Cu, Fe, Mn, Ni, Pb, and Zn) in the samples.

Throughout, Fe, Mn, and Zn displayed the highest concentrations in water samples for Bindal (Fe: 0.97 - 4.9 mg/L; Mn: 0.1-1.6; Zn: 0.11 - 2.9) and Rispana (Fe: 0.7 - 38; Mn: 0.09 - 4.9; Zn: 0.3 - 7.3; SI 2&3). Similarly, in sediment samples, these heavy metals were most prevalent in Bindal (Fe: 20 - 4273.61 mg/L, Mn: 2 - 68.6, Zn: 0.6 - 61.4) and Rispana (Fe: 16 - 8854.12 mg/L, Mn: 0.5 - 308.7, Zn: 0.03 - 46.5; SI 2 and SI 3). Still, the remaining heavy metals were also detected throughout, reaching maximum concentrations of 28.83 Ag μg/L, 98.3 Cd μg/L, 24.84 Co μg/L, 485 Cr μg/L, 122.2 Cu μg/L, 331.24 Ni μg/L, and 1073.8 Pb μg/L in Bindal waters, 1.37 Ag mg/L, 2.2 Cd mg/L, 3.1 Co mg/L, 5.2 Cr mg/L, 20.3 Cu mg/L, 7.3 Ni mg/L, and 11.4 Pb mg/L in Bindal sediments (SI 2 and SI 3). Similarly, Ag, Cd, Co, Cr, Cu, Ni, and Pb were detected in Rispana water and sediment samples at maximum concentrations of 73.77, 72.42, 44.27, 1542, 1775, 657.12, and 1272.58 μg/L in water and 1.8, 0.22, 5.1, 9.8, 23.03, 14.22, and 18.21 mg/L in sediment samples, respectively (SI 2 and SI 3). Unlike fecal indicators, the maximum concentrations of heavy metals in both water and sediment samples across seasons were detected often in the sampling locations downstream of the WWTPs for each river however they weren’t significantly different from other sites (all p > 0.05, Kruskall-Wallis test; SI 2 and SI 3) indicating a substantial contribution of wastewater. Unsurprisingly, heavy metal concentrations were usually significantly higher in sediment samples compared to water samples in Bindal (summer: p = 0.0074 - 0.0097, Wilcoxon-test, monsoon: p = 0.011 - 0.04 and, winter: p = 0.0079 - 0.0097) and Rispana (summer: p = 0.0086 - 0.025, Wilcoxon-test, monsoon: p = 0.002 - 0.02 and, winter: p = 0.0021 - 0.0047) with only a few exceptions in Bindal monsoon (Cd, Cr, Ni, Pb) and Rispana summer samples (Cr, Mn, Ni, Zn) (SI 2 and SI 3).

Unlike fecal pollutants, heavy metal concentrations generally displayed no significant seasonal variation in concentration for water and sediment samples in both Bindal and Rispana (all p > 0.05, Wilcoxon signed-rank test; Fig. 4). Here the lone exceptions were Pb concentrations, which were significantly higher in monsoon than in summer in Rispana water (p = 0.012, Wilcoxon signed-rank test) and Co, Cu and Ni, which were significantly higher in winter than in summer in Rispana sediments (p = 0.012, Wilcoxon signed-rank test; Fig. 4).

**Fig. 4.**
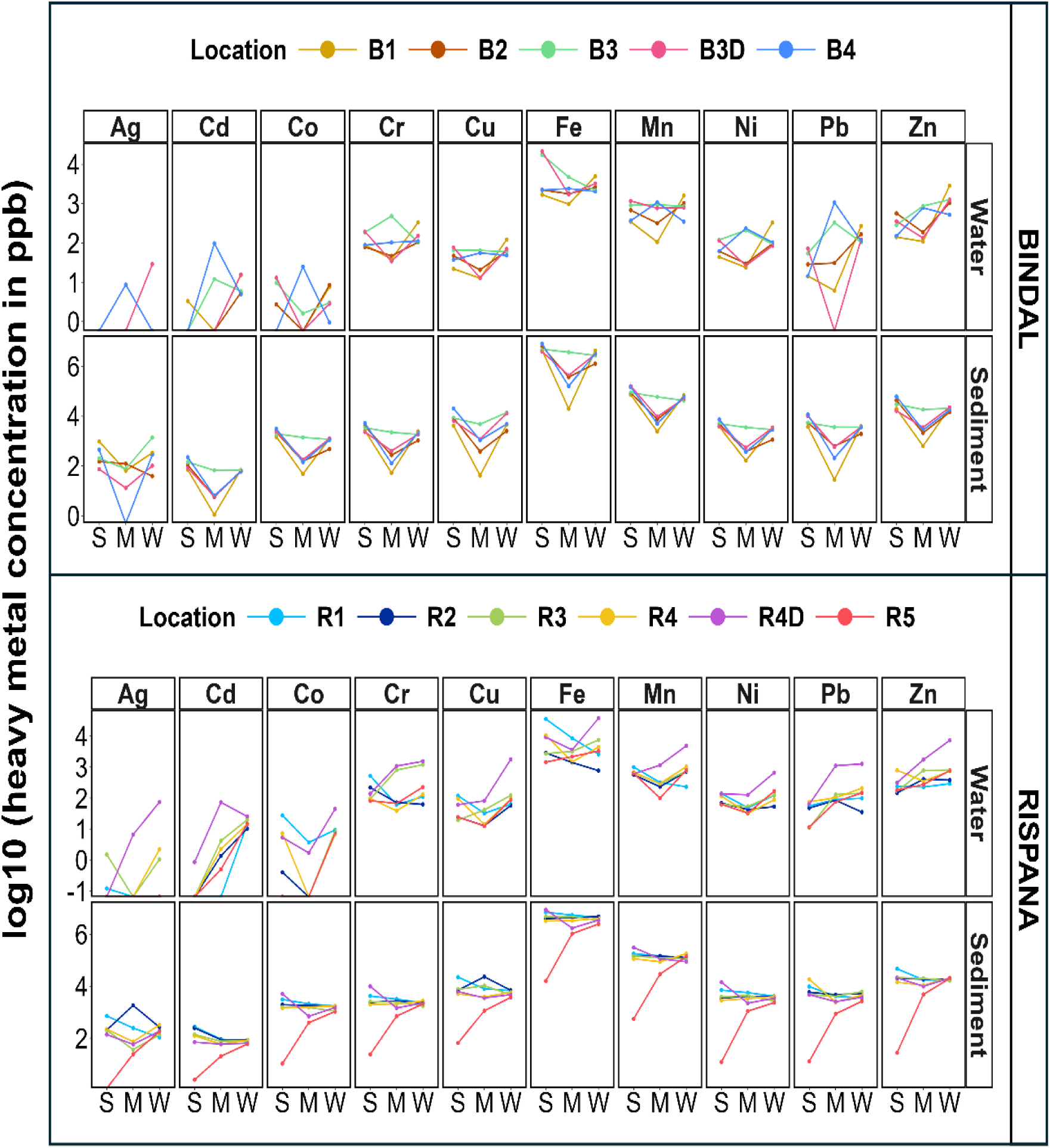
Variation in heavy metal concentrations in water and sediment samples for Bindal and Rispana during summer, monsoon and winter.

In conclusion, high concentrations of diverse heavy metal pollution were detected in each river across seasons, with concentrations significantly higher in sediments than in water samples. However, heavy metal pollution displayed no significant seasonality, as observed for fecal pollution.

### Identifying drivers of the spatiotemporal distribution of ARGs in Bindal and Rispana rivers

Since both, fecal indicators and co-selective agents were detected at high prevalence for both the urban rivers, we evaluated the relative abundance of model ARGs *erm*F, *sul*1, *sul*2, and *tet*W. Throughout, high levels of ARG pollution were detected in both rivers, with average relative abundances of the ARGs ranging between 2.3×10^-1^ and 2.6×10^-3^ in water and between 9.9×10^-1^ and 1.1×10^-1^ in the sediment samples (SI 4 and SI 5).

For water samples, no significant changes in the relative abundance of *erm*F, *sul*1, *sul*2, and *tet*W were detected along the flow direction of Bindal in summer (p = 0.18 - 0.45, Spearman rank correlation) and winter (p = 0.18 - 0.25, Spearman correlation), while in monsoon a slight but significant decrease for ARGs *erm*F (p = 0.027) and *tet*W (p = 0.027) were observed (SI 4). Similarly, no significant difference in relative abundance along the length of Rispana was observed for any season (p = 0.56 - 0.834, Spearman correlation) with the lone exception of increase in *erm*F with flow direction in winter (p < 0.0001; SI 4). For sediments, no significant differences in the relative ARG abundance were observed for either river or season (p > 0.05), with the lone significant exception being an increase in *erm*F abundance over the length of Rispana during monsoon (p < 0.01; SI 5).

ARG relative abundances followed strong seasonality patterns: The relative abundance of *erm*F, *sul*2, and *tet*W across locations was consistently significantly elevated by up to 2-3 orders of magnitude in monsoon compared to summer for both Bindal and Rispana water samples (*erm*F: p = 0.032; *sul*2: p = 0.00097; *tet*W: p = 0.0068; Wilcoxon signed-rank test; Fig. 5). While similar trends were observed for sul1 at certain locations, especially of Rispana, this was not consistent across locations (p = 0.123). After monsoon, the relative abundance of all ARGs consistently and significantly dropped in winter (*erm*F: p = 0.004; *sul*1: p = 0.024; *sul*2: p = 0.00097; *tet*W: p = 0.00195; Fig. 5) by up to 2 orders of magnitude. There was no consistency in return of the ARG abundances after monsoon (winter) to pre-monsoon (summer) levels: *sul*2 remained up to 1 order of magnitude higher compared to summer samples (p = 0.00976; Fig. 5), while *erm*F and *sul*1 levels significantly decreased by up to one order of magnitude (*erm*F: p = 0.0.004; *sul*1: p = 0.00097) and no significant difference was observed for *tet*W between winter and summer (p = 0.053). Similarly, in sediment samples, the relative abundance of ARGs was consistently significantly higher by between 0.4 and 1.5 orders of magnitude in monsoon compared to pre-monsoon summer samples at most locations for both Bindal and Rispana (all p < 0.05; Fig. 5). However, ARG abundance slightly decreased for most sediment samples between monsoon and winter, except for *sul*1 (p = 0.032). This also resulted in post-monsoon winter sediment samples having significantly elevated relative ARG abundances compared to post-monsoon summer (all p < 0.05; Fig. 5) by up to 1 order of magnitude.

**Fig. 5.**
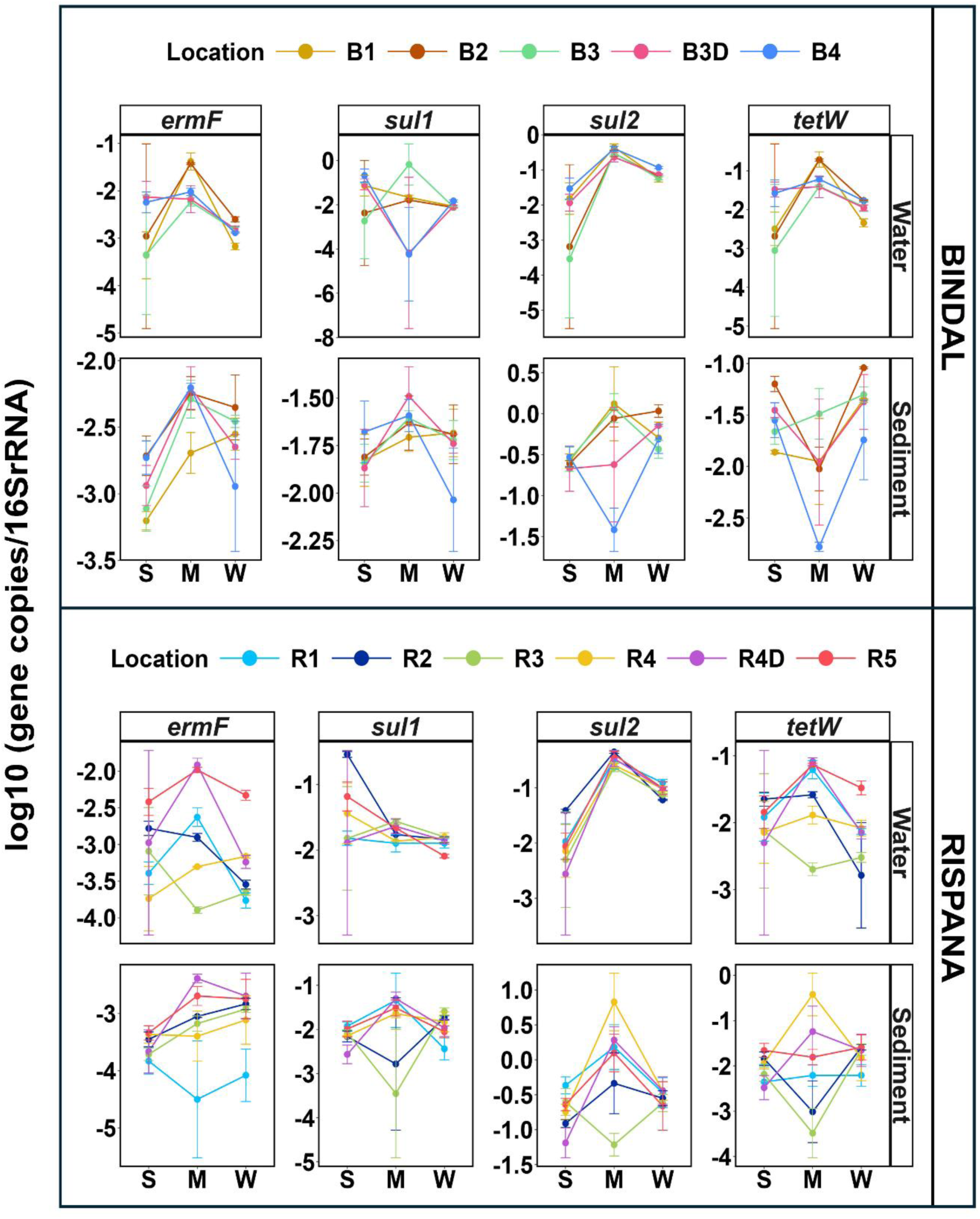
Relative abundance of ARGs in water and sediment samples from Bindal and Rispana during summer, monsoon and winter. The points are the mean (n =3) with standard deviation in the log_10_ graph.

To identify if the high levels of observed ARGs originated from fecal pollution, due to co-selection through heavy metals or both, we evaluated the correlation between the abundance of ARGs (*erm*F, *sul*1, *sul*2 and *tet*W) with fecal indicators (*E. coli* and *int*I1) and heavy metal concentrations in water and sediment samples of both rivers. Throughout, all ARGs positively correlated with the fecal indicator, *int*I1, across different seasons for water and sediment samples in both rivers (all p < 0.001, R > 0.7, Spearman rank correlation) and with *E. coli* in the case of water samples, when it was above the limit of detection (all p < 0.001, R > 0.73; Fig. 6). However, no correlation between fecal indicators and heavy metals or ARG and heavy metals were detected in water samples (p > 0.05; Fig. 6). Similarly, in sediment samples, most heavy metals did not correlate with either fecal indicators or ARGs (all p > 0.05, Spearman rank correlation), with the exception of Rispana sediments, where Co, Cu, and Ni weakly and positively correlated with *sul*2 (R < 0.35, p < 0.05) and Ag, Cd, Co and Cu positively correlated with fecal indicator *int*I1 (R < 0.3, p < 0.05; Fig. 6).

**Fig. 6.**
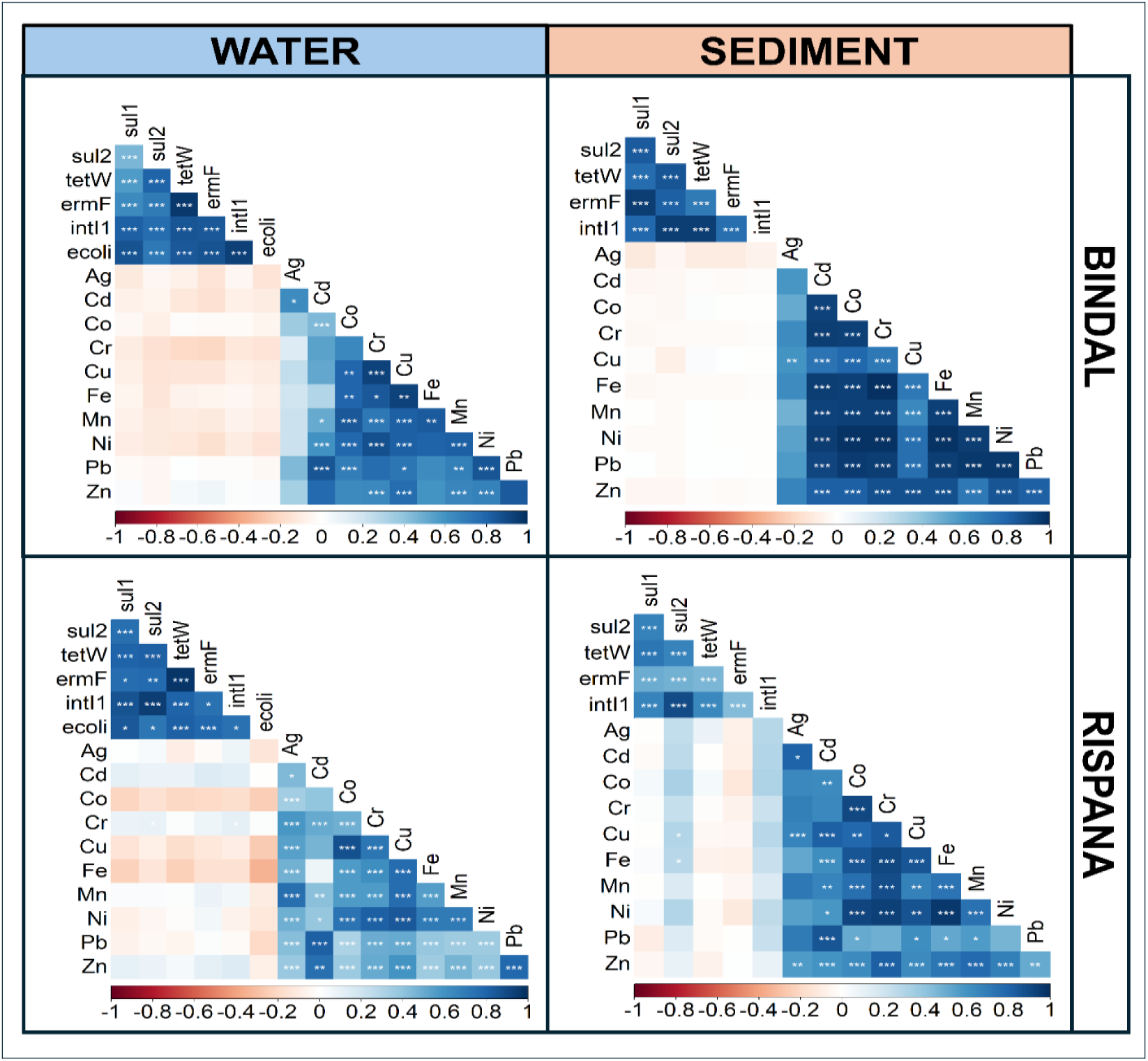
Spearman rank correlation between the absolute ARGs, absolute fecal indicators and heavy metals for water and sediment samples in all seasons (summer, monsoon, winter) combined.

Overall, both rivers in all three seasons contained high levels of ARGs with no significant trend in the direction of the flow of the rivers. Regarding seasonality, ARG abundance increased significantly from the summer to the monsoon season and significantly decreased back to pre-monsoon levels in winter in water samples. However, ARG levels remained elevated in post-monsoon winter compared to pre-monsoon summer samples. Fecal pollution was identified as the main driver underlying the spatiotemporal patterns of ARGs in Indian urban rivers, while heavy metal pollution and the resulting co-selective pressures displayed no significant effect.

## Discussion

We aimed to determine whether the river resistome receiving untreated wastewater is driven by fecal contamination or co-selective agents such as heavy metals and how these dynamics are affected by heavy rainfall during the monsoon season. The presence of fecal indicators and ARGs in the rivers were elevated right after entering the city area and showed no significant variations along their flow paths within the city, indicating heavy pollution due to the discharge of untreated wastewater along the entire length of the rivers. The fecal indicators and ARGs were relatively higher in monsoon, suggesting that rainfall brings a high load of fecal contaminants via contaminated surface run-off, including resistant bacteria, into the rivers. Despite the heavy metals being present at high concentrations, there was no evidence of a significant contribution of co-selection to the observed ARG dynamics in either river.

The discharge of wastewater into both rivers is evidenced by elevated total bacterial load and the presence of fecal indicators, *E. coli* and *int*I1, which are commonly associated with wastewater contamination [7], [28], [37]. The observed levels of absolute fecal indicators along the length in both rivers can be explained by the incoming load of total bacteria since the 16S rRNA gene copies positively correlated with the absolute values of fecal indicators for water and sediment samples. Moreover, the positive correlation between the abundance of *E. coli* and *int*I1 at every location for both rivers suggests that the *E. coli* detected in samples was likely from fecal contamination. Unlike other studies reporting high levels of resistant *E. coli* in Indian rivers [19], [25], [38], *E. coli* was below detection limits in sediment samples from this study. Potential reasons include competition with native microbes, predation, unfavorable conditions [39], [40], or insufficient DNA yield due to a single extraction cycle [41]. Thus, *int*I1 may be a more reliable indicator of fecal pollution in this context.

The strong and positive correlation observed between ARGs - *erm*F, *sul*1, *sul*2, and *tet*W and fecal indicators suggests that both share a common source of origin. This implies that the ARGs detected in the river are likely introduced through the same pathway as fecal contamination, further supporting the notion that wastewater is the primary contributor to the presence of both fecal indicators in the river system. The ARGs targeted in this study are well known to be associated with wastewater and, hence, are a strong marker for wastewater discharge into the river [9], [42], [43], [44].

Contrary to the usual trend where ARG abundance and fecal indicators increase downstream due to cumulative pollution [20], [45], [46], [47], there was no significant spatial variation in the relative abundance of fecal indicators or ARGs at different time points. This suggests that the wastewater is not discharged from a single point source but rather distributed throughout the river system. Given that the 18.2 and 9.4 MLD WWTPs are situated downstream of Bindal and Rispana, the observed levels of wastewater indicators upstream are likely attributable to the discharge of untreated wastewater through various drains [26].

The effect of rainfall is prominent in water and sediment samples for both rivers, as the relative abundance of fecal indicators and ARGs increased significantly during the monsoon season compared to summer (Fig. 3 and 6). The findings are similar to some reported cases where the effect of rainfall increases the ARG abundance [48], [49], [50], while some studies showed a lower abundance of ARGs in rivers in the wet season as a result of dilution[45], [51]. One of the reasons for the elevated level of ARGs is reported to be co-selection by heavy metals present in the river [12], [22], [24] however, in this study, the trends observed in season-wise heavy metal concentrations are not the same as the ARGs. The heavy metals do not show any significant difference between seasons, and the rainfall didn’t affect their concentration level. Despite their presence in high concentration, the heavy metals analysed in water and sediment samples don’t correlate with the ARGs. The MCSC (minimum co-selective concentration) suggested by Seiler and Berendonk Cd, Cu, Ni, Pb and Zn for river surface water is as low as 0.03 ug/L, 1.5 ug/L, 0.29 ug/L, 0.15 ug/L, and 19.61 ug/L, respectively [52]. The heavy metal concentrations in this study were mainly above the suggested MCSC for both rivers. One possible reason could be that a fixed limit for minimum inhibitory concentration (MIC) and MCSC is not applicable in general since it depends on multiple factors, including pH, environmental matrix, and sensitive toxicity level [53]. Moreover, the absence of a correlation between heavy metals and ARGs, as well as between heavy metals and fecal indicators, suggests that the sources responsible for the biological and chemical contamination are distinct (Fig 8). Typically, wastewater comprises black water from toilets/septic tanks and grey water from kitchens and washrooms, eventually conveyed to the WWTP via a sewer line. Black water has biological contaminants like pathogens and ARGs, while grey water has chemical pollutants like PCPs, heavy metals and disinfectants [6], [54], [55], [56]. The higher level of ARGs and ARBs in the river during heavy rainfall could be the transportation of effluent from septic tanks which aren’t connected to the sewer lines. India, like many LMICs, depends on septic tanks since the WWTPs do not cover most cities and sewer lines are limited [54], [56]. Without a sewer line, grey water is discharged through open drains into the nearest water body. In contrast, black water is allowed to percolate underground, seeping into the soil, groundwater and surface water. The effluent with a higher level of a plethora of pathogens and ARGs from overflowing septic tanks is known to seep through the ground and contaminate the surface waters during heavy rainfall [56], [57], [58].

Other than the effect of precipitation, the seasons were expected to affect the trends in the levels of ARGs in the river water. The abundance of *sul*2 was significantly higher in winter than in summer for water samples and was more prevalent in sediment samples, where all the ARGs (except for *sul*1) and *int*I1 were significantly higher in winter than in summer. A few studies reported a higher abundance of ARGs in summer than in winter, especially when the water bodies were glacier-fed, which contradicts the observations in this study, in which both rivers rely on the urban drains for a substantial portion of their flow [47], [59]. Sabri et al. (2020) reported certain genes were higher in winter (*erm*B), some lower (*sul*1, *sul*2, and *int*I1) and had no change (*tet*W) relative to summer [46]. In the case of this study, the *erm*F and *sul*1 in water were higher during the summer compared to winter, and *tet*W and *int*I1 remained similar in both seasons. Seasonal temperature is linked with increased total ARG abundance by enriching the biomasses [60]. The presence of genes conferring resistance to a particular class of antibiotics in wastewater has been associated with certain seasons, likely due to the use of antibiotics relevant to that season [30], [46], [61]. A study observed that the level of ARG abundance significantly rises within the microbial community during a rainfall event; however, it eventually falls back to its original level before rainfall [62]. In this study, the monsoon did have elevated levels of ARGs that did not return to the pre-monsoon levels, especially in sediment samples (Fig. 5). The elevated levels of ARGs in sediment could be because sediment is better at harbouring and protecting incoming extracellular DNA [63].

Two ARGs - *erm*F and *sul*2, were consistently higher in monsoon than in summer in both water and sediment samples. Many studies reported *sul*2 higher than *sul*1 in the samples [64]. While one study reported no significant seasonal change in the level of *sul*1, *sul2* showed a significant change in spring [65]. Unlike *sul*1, *sul*2 is highly proliferated in environmental settings due to its occurrence in multiple transmissible and small non-conjugational plasmids [66]. Pei et al. 2006, reported an increase in the prevalence of the *sul*2 gene abundance with high urban and agricultural activity while being undetectable in relatively pristine sites [44]. In contrast, *sul*1 was ubiquitous in upstream and downstream locations, suggesting that *sul*2 may serve as a more sensitive indicator of anthropogenic pollution. This finding is consistent with the observation in this study. The ARG *erm*F has been reported to be unique to Asian and American wastewater and not detected in European countries [67].

## Conclusion

We demonstrate that the main source of the resistome in urban rivers, Bindal and Rispana, is the constant inflow of untreated wastewater throughout the year and not the co-selection of ARGs due to the presence of heavy metals. This may be true for many Indian rivers that receive untreated or partially treated wastewater, solid waste, and greywater via urban drains. The rainfall adds to the ARG abundance of the river instead of diluting it. In such cases, targeting the key fecal sources, such as septic tanks and managing their overflow during monsoon season can help manage the source better. This can include adding a third compartment and unloading the tank pre-monsoon every year.

The MGE *int*I1 is more reliable than *E. coli* as an indicator for fecal pollution and horizontal gene transfer. Heavy metals, despite being present in the river in higher concentrations than the MCSC, were not linked with the co-selection of ARGs in river water and sediments.

## Acknowledgements

The authors thank Deepika Bhaskar, Saurabh Dixit, and Ojash Giri for their help with heavy metal analysis and qPCR standard preparation.

## Supplementary information

**SI 1.**
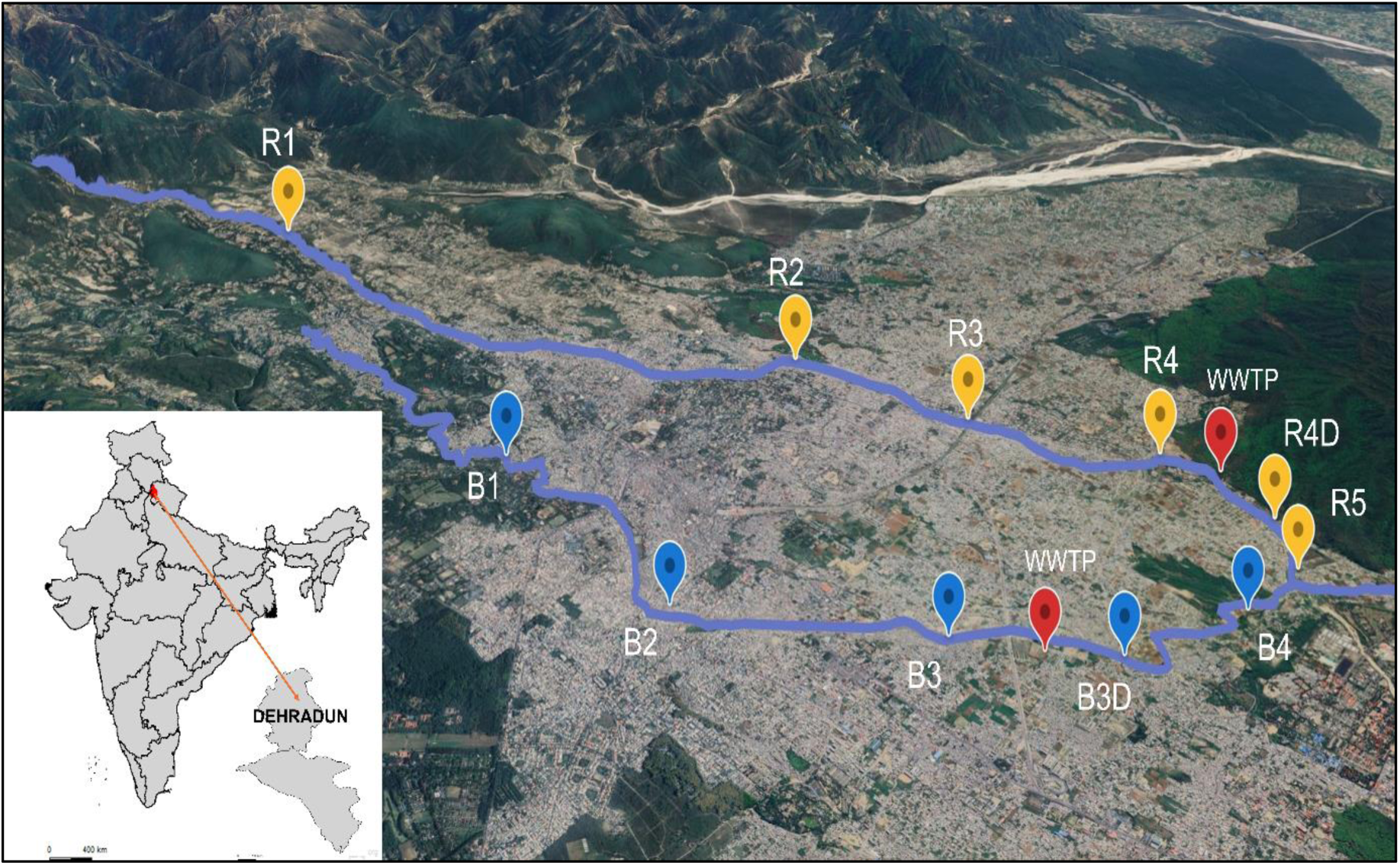
Different sampling locations on the banks of the Bindal and Rispana with two WWTPs downstream of the river highlighted with red pushpins on the map. The two sub-tributaries confluence together to become the river Suswa, which merges with river Song before combining with the river Ganga (Maps: Google earth web; GADM.org).

**SI 2.**
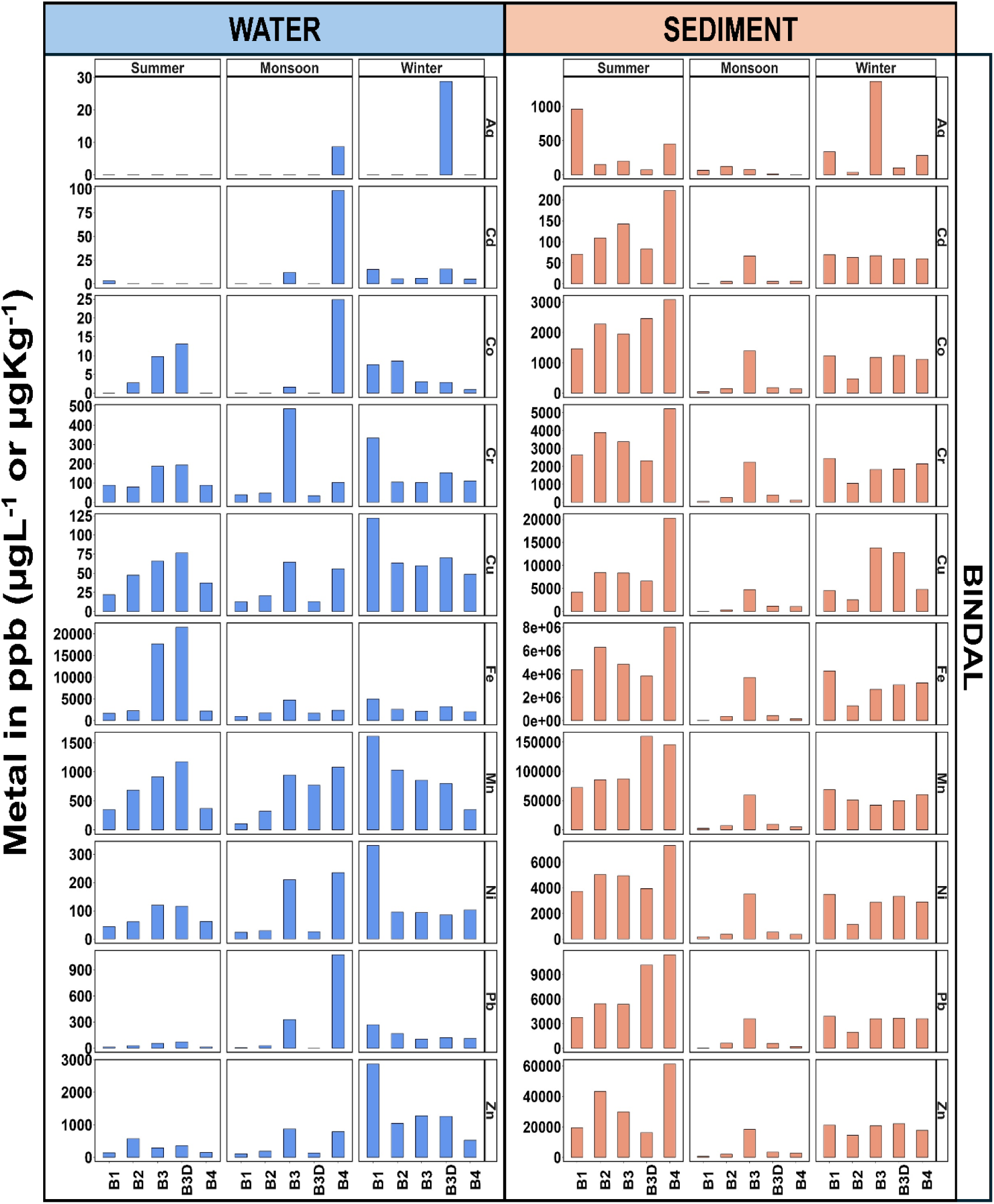
Heavy metal concentrations (ug L^-1^ or ug Kg^-1^) in water and sediment samples at different Bindal and Rispana sampling locations during summer, monsoon and winter.

**SI 3.**
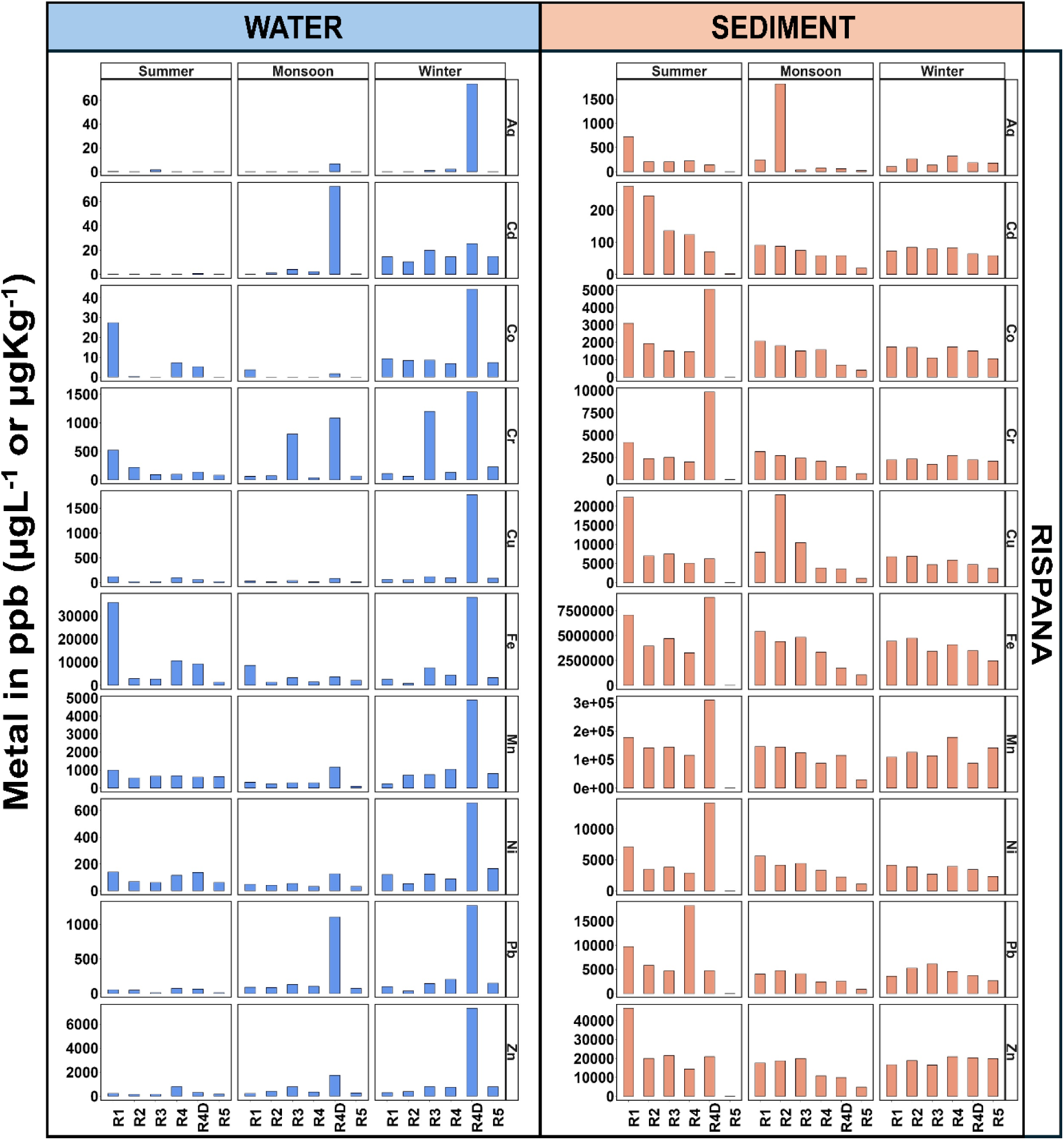
Heavy metal concentrations (ug L^-1^ or ug Kg^-1^) in water and sediment samples at different Bindal and Rispana sampling locations during summer, monsoon and winter.

**SI 4.**
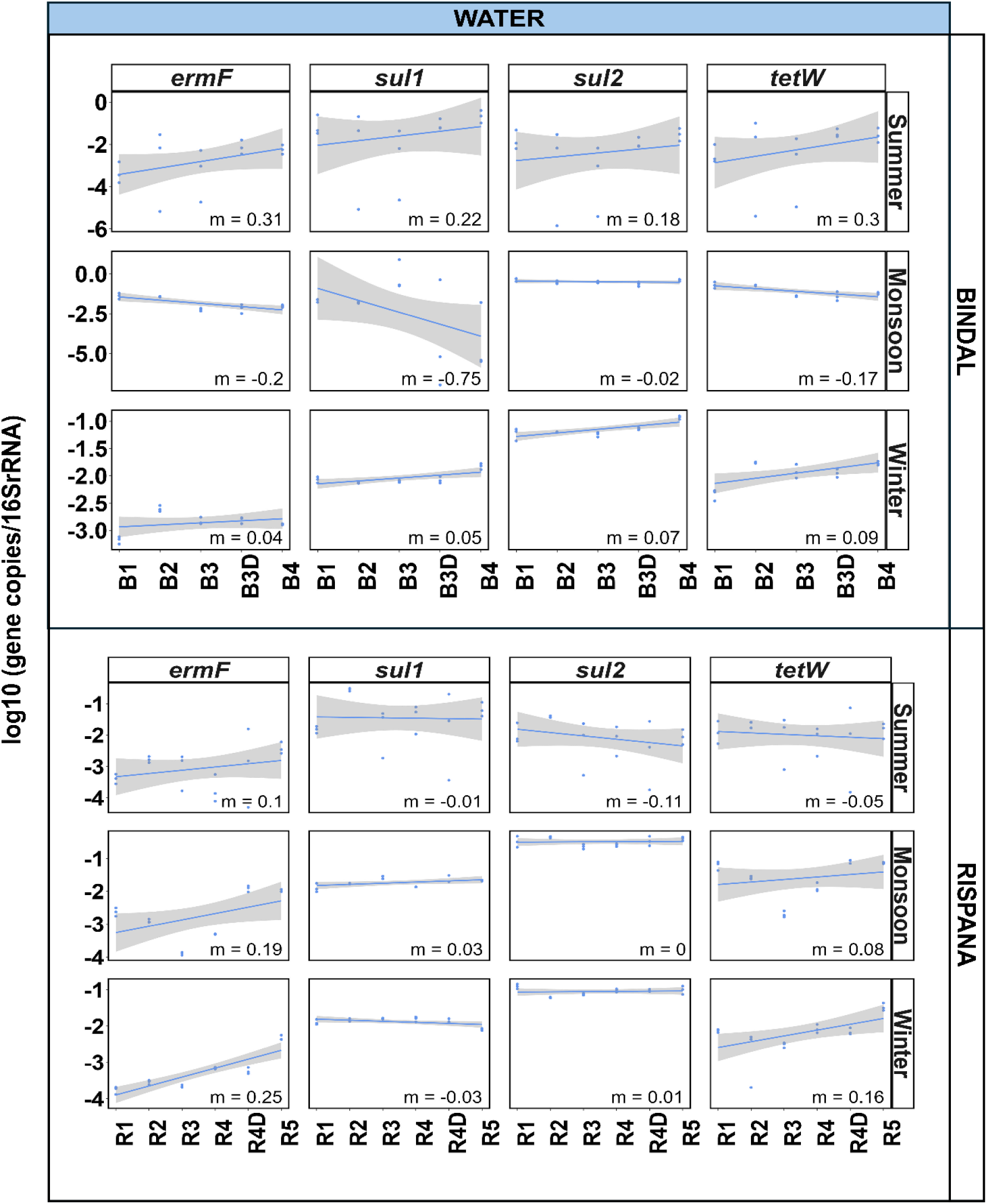
Temporal dynamics of the ARGs (*erm*F, *sul*1, *sul*2, and *tet*W). Linear regression for ARGs normalized to 16S rRNA along the length of Bindal and Rispana for water samples in summer, monsoon and winter.

**SI 5.**
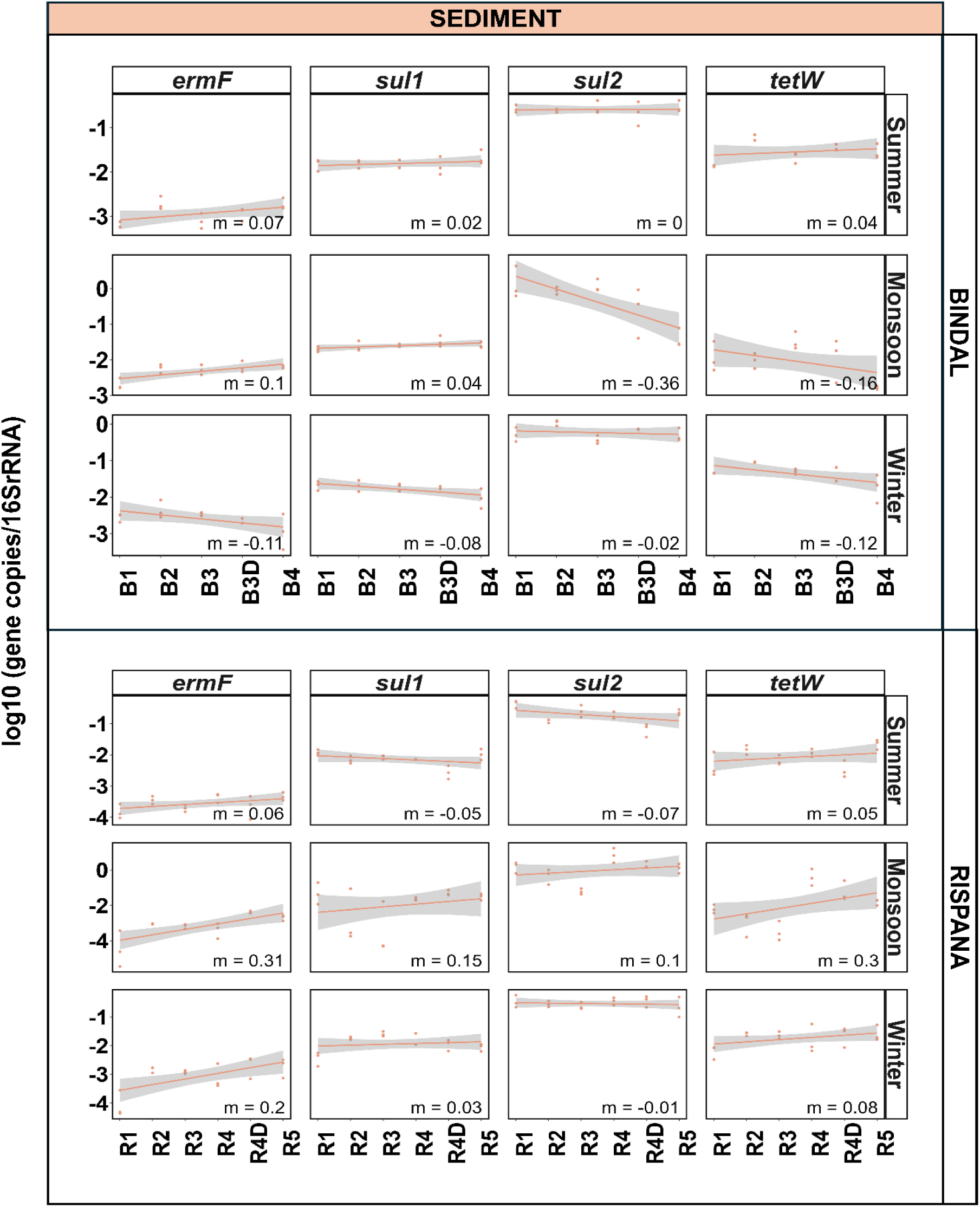
Temporal dynamics of the ARGs (*erm*F, *sul*1, *sul*2, and *tet*W). Linear regression for ARGs normalized to 16S rRNA along the length of Bindal and Rispana for sediment samples in summer, monsoon and winter.

